# Edge effect on the distribution of the Green Shield Bug *Palomena prasina* in hazelnut orchards, and the role of adjacent habitats in crop colonization

**DOI:** 10.64898/2026.01.08.694378

**Authors:** Laetitia Driss, Christophe Andalo, Chloé Liedtke, Rachid Hamidi, Alexandra Magro

**Affiliations:** Association Nationale des Producteurs de Noisettes (ANPN), 1500 route de Monbahus, Cancon, France; Centre de Recherche sur la Biodiversité et l’Environnement (CRBE), Université de Toulouse, CNRS, IRD, Toulouse INP, Université Toulouse 3 – Paul Sabatier (UT3), Toulouse, France; Université de Toulouse - ENSFEA, 2 rt de Narbonne, 31326 Castanet-Tolosan, France

**Keywords:** Host plants, IPM, spatio-temporal distribution

## Abstract

*Palomena prasina* (L.) (Hemiptera: Pentatomidae), the green shield bug (GSB), is an important hazelnut pest in Southern Europe. Currently, its control focuses on insecticide spraying during the crop season. We hypothesised that, as for other pentatomid species, adjacent habitats strongly influence the population build-up in spring and, therefore, lead to edge effects in crop fields during the production season. This could allow for precision-targeted pest management strategies. This study examined the spatio-temporal dynamics of the GSB in spring and summer over two years. We investigated the plant preferences of GSBs in spring and the existence of edge effects on their distribution and on fruit damage in hazelnut orchards in summer. We also examined the relative contribution of adjacent habitats to GSB abundance in crops.

Our results show that, following their emergence from overwintering in spring, GSB adults, and later on their offspring, are primarily found on wild host plants in natural habitats, particularly on *Crataegus monogyna* Jacquin, and *Cornus sanguinea* L.. In early summer, the older nymphs of the second generation colonise the hazelnut orchards, with populations and damage proportion displaying edge effects. We were unable to identify an adjacent habitat variable that significantly explains the abundance of GSB in the orchards, although both forest habitats and hazelnut orchards have an effect on GSB abundance. Based on these findings, we propose various IPM strategies for controlling *P. prasina*.

## Introduction

The Pentatomidae family (Hemiptera), commonly known as stink bugs, is one of the largest heteropteran families (Grazia *et al*., 2015; Panizzi & Lucini, 2017). Most species in this family are highly polyphagous (Panizzi, 1997): for example, the brown marmorated stink bug *Halyomorpha halys* Stal can feed on 88 plant species (Bergmann *et al*., 2016), whereas the green stink bug *Nezara viridula* (L.) can feed on plant species belonging to 30 different families (Panizzi, 1997). Stink bugs feed on both wild and crop plants, and many species are therefore economically important agricultural pests. Indeed, the feeding punctures caused by stink bugs lead to necrosed or distorted fruits (Acebes-Doria *et al*., 2016a; Chocorosqui & Panizzi, 2004; Cissel *et al*., 2015), blank nuts (Hedström *et al*., 2014) and negatively affect growth and productivity (Huang *et al*., 2014). Consequently, they result in significant economic losses to crops (Rice *et al*., 2014; Esquivel *et al*., 2018).

The control of stink bugs commonly involves the use of broad-spectrum insecticides, such as Neonicotinoids or Pyrethroids (Saruhan *et al*., 2023; Wiman *et al*., 2023). However, the use of some of these pesticides is restricted in several countries due to their impact on human health and the environment (Kathage *et al*., 2018; Ravula & Yenugu, 2021), secondary pests outbreaks (Gerson & Cohen, 1989), and pest resistance (Sosa-Gómez *et al*., 2020). Therefore, it is paramount to look for more sustainable alternatives for controlling stink bugs, in a perspective of Integrated Pest Management (IPM), with pesticides being applied to crops only as a last resort. In this context, precise information on the temporal and spatial dynamics of pests is crucial.

Pest outbreaks are often unevenly distributed, but early detection and precision-targeted pest management can limit the need for pesticide spraying (Iost Filho *et al*., 2020). Edge-biased distributions have been commonly observed in the spatial distribution of insects, including in transition zones between natural habitats and cultivated areas (Nguyen & Nansen, 2018). For example, *Diaphorina citri* Kuwayama (Hemiptera: Liviidae), an economically important citrus pest, and *Epitrix tuberis* Gentner (Coleoptera: Chrysomelidae), a potato pest, exhibit strong field edge effects (Sétamou & Bartels, 2015; Vernon & Van Herk, 2017); in both cases, control measures focused on the edges of crop fields are recommended (Vernon & Van Herk, 2017; Miranda *et al*., 2021). Edge effects have also been observed for stink bugs, such as for *H. halys* in soybean fields, where higher abundances and damage rates are observed at the edges of fields, which leads to targeted chemical control, resulting in 85-95 % reduction in pesticide spraying (Leskey *et al*., 2012; Aigner *et al*. 2017).

Although identifying and characterising edge effects offers undeniable opportunities to control pests once they colonise the crops, understanding the source of the dispersing populations and their dispersal dynamics is also important. During the harsh season, stink bugs overwinter in the leaf litter, under the bark of trees, or in urban structures (Driss *et al*., 2023). Once they resume development, post-overwintering adults start feeding and reproducing on herbaceous plants, shrubs, or trees (Panizzi & Lucini, 2017), moving between host plants according to their preferred phenological stages (McPherson & McPherson, 2000), which is primordial for their fecundity and survival (Panizzi, 1997). Extensive monocultures of crop plants at the preferred phenological stage for stink bugs, are highly attractive as they represent a large food resource. In this context, identifying host plants and, more broadly, the spatial dynamics of stink bugs between habitats can help to manage their populations (Aigner *et al.,* 2017).

The green shield bug (GSB), *Palomena prasina* (L.) (Hemiptera: Pentatomidae), is a polyphagous stink bug that is widely distributed across the Palearctic region (Lupoli & Dusoulier, 2015). In hazelnuts, feeding punctures lead to empty or aborted fruits and shrivelled or necrosed kernels (Romero *et al*., 2009; Tuncer *et al*., 2005; Tavella *et al*., 2001). GSB is now considered a major pest in European commercial hazelnut orchards (Tuncer *et al*., 2005; Ak *et al*., 2018; Hamidi *et al*., 2023). Currently, GSB control mostly involves insecticide spraying at the beginning of summer, but this has had limited success (Bayle, 2022; Hamidi *et al*., 2023), making alternative control measures urgently needed.

According to Saruhan *et al*. (2023), GSB is univoltine. Driss *et al*. (2023) demonstrated that the adults overwinter in leaf litter in various ecosystems, including hazelnut orchards. While it cannot be ruled out that post-overwintering adults remain in the orchards, it should be noted that they have been observed feeding on various herbaceous and woody plants alongside their offspring (Boselli, 1932; Javahery, 1967). Furthermore, GSB has mainly been seen in the crop from the end of spring (Hamidi *et al*., 2022). Therefore, we hypothesize that, at the end of the winter, GSB moves from the hazelnut orchards to adjacent habitats where it finds other host plants during the fruiting period. It then returns to the orchards when the hazelnuts are present, ultimately resulting in an edge effect distribution in the crop. Furthermore, we anticipate that among the various adjacent habitats, diverse natural habitats should prevail to elucidate the crop summer abundance of GSB, in contrast to certain overwintering habitats such as the hazelnut orchards.

The aim of this study was therefore threefold a) to identify if, in spring-summer, GSB is present in different host plants, including hazelnut trees, and to evaluate its preferences, including plant phenological stage, using relative abundance as a proxy, b) to investigate whether there is an edge effect on hazelnut orchards during the crop production season with regard to the distribution of GSB individuals and damage, and finally c) to test which adjacent habitats better explain the abundance of GSB in hazelnut orchards. The results will be discussed in terms of possible IPM strategies.

## Material and methods

### 1.1 Host plant preferences

#### Study sites

Three study sites, spaced 6 km to 11 km apart, were selected near Cancon, in France (44.539383N, 0.610745E). At each site, the study area comprised one hazelnut orchard and the adjacent diverse wild hedges facing the orchard. Two of the orchards were of the Corabel cultivar and one was of the Pauetet cultivar. Field surveys were conducted once a week on the same day at all sites between March and mid-August in both 2021 and 2022.

#### Insect sampling on host plants

At each site, GSB was prospected in hazelnut trees planted at the edge of the orchard and in nine plant species commonly found in hedgerows in southwestern France (Badeau *et al.,* 2017) (see Table 1). Ten individuals were sampled for each species. For each individual, two main branches located at approximately eye level were scouted above a beating tray (116 cm x 86 cm) to collect GSBs. Furthermore, a 25-metre-long, 1-metre-wide grass strip containing a variety of species (Table 1), located between the edge of the hazelnut orchard and the hedge, was also sampled using a sweep net. Twenty sweeps were performed along the strip. All the collected GSBs were brought to the laboratory, where the different developmental stages were identified and counted. The presence of the first egg masses was noted.

**Table 1.**
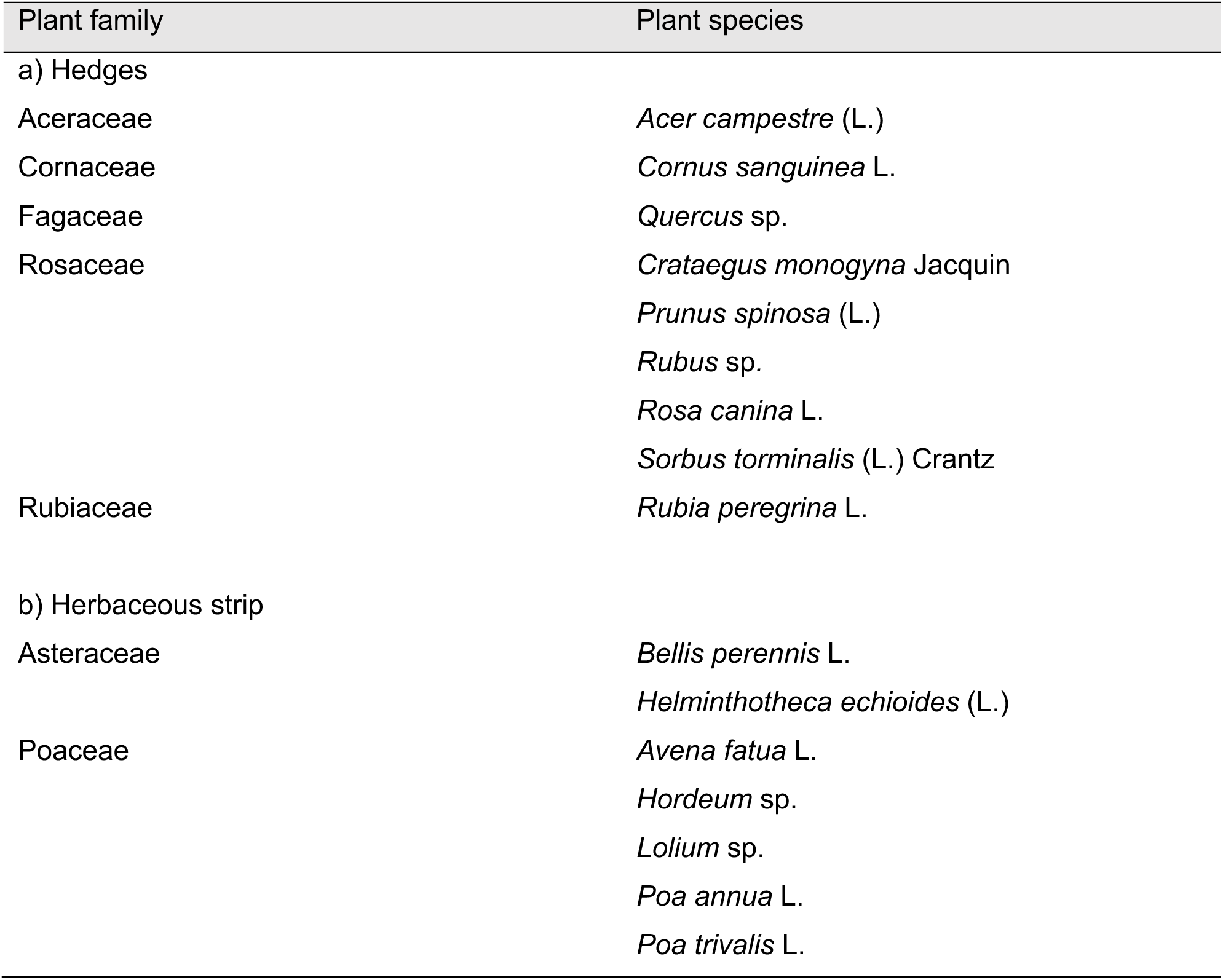
List of wild plant species monitored in 2021 and 2022 in a) the hedges, and b) in the herbaceous strip surrounding the hazelnut orchards.

| Plant family | Plant species |
| --- | --- |
| a) Hedges |  |
| Aceraceae | *Acer campestre* (L.) |
| Cornaceae | *Cornus sanguinea* L. |
| Fagaceae | *Quercus* sp. |
| Rosaceae | *Crataegus monogyna* Jacquin |
|  | *Prunus spinosa* (L.) |
|  | *Rubus* sp. |
|  | *Rosa canina* L. |
|  | *Sorbus torminalis* (L.) Crantz |
| Rubiaceae | *Rubia peregrina* L. |
| b) Herbaceous strip |  |
| Asteraceae | *Bellis perennis* L. |
|  | *Helminthotheca echinoides* (L.) |
| Poaceae | *Avena fatua* L. |
|  | *Hordeum* sp. |
|  | *Lolium* sp. |
|  | *Poa annua* L. |
|  | *Poa trivialis* L. |

Additionally, the phenology of the scouted plants in the hedge and orchard was recorded weekly for each individual plant according to three categories adapted from Badeau *et al.,* (2017): (1) dormancy, (2) presence of flowers (at least 10 % of flowers in the “bud” to “bloom” stage), (3) presence of fruits (at least 10 % of fruits in the “unripe” to “mature” stage).

### 1.2 GSB spatial distribution in hazelnut orchards

#### Studied orchards

Due to the unavailability of the orchards sampled during the host preference study, alternative orchards were selected in the same area. Each year, two old hazelnut orchards (planted in 1980 and 1997, with trees 3–5 meters high) and two young hazelnut orchards (planted in 2018 and 2019, with trees 2–2.5 meters high) were chosen. In 2022, one of the young orchards was replaced by another. None of the orchards were sprayed with insecticide during the study. The vegetation within the hazelnut orchards was mown regularly, except in one of the old orchards. The hazelnut trees were grown in rows, 5 meters apart and 5 or 2.5 meters apart within a row, respectively for the old and young orchards.

#### Insect sampling in the hazelnut orchards

GSB monitoring took place once a week in both 2021 and 2022.

In 2022, monitoring took place from mid-June (week 25) to mid-August (week 32). In each orchard, four opposite sides were selected, and sampling was performed in a zone corresponding to five rows of hazelnut trees, at distances of 0, 5, 10, 20 and 30 meters from the orchard edge. In old orchards, ten consecutive trees were selected in each row, with two branches (at approximately eye level) of each tree being scouted. In young orchards, one branch (at approximately eye level) of twenty consecutive trees in each row was scouted. The branches were sampled using a beating tray as described above. All the collected GSB were brought to the laboratory, where the different developmental stages were identified and counted.

In 2021, the GSB monitoring followed the same design as in 2022, but due to logistical constraints there were two exceptions. The monitoring started one week later, and, during the first two weeks, the trees located 30 meters from the edge were not sampled. Consequently, the GSB abundance data collected in 2021 was only used in the analyses described in section 1.3 (adjacent habitats).

#### Assessment of damage

The damage rate was assessed based on the number of hazelnuts collected from the orchards during the 2022 harvest season (i.e. from late August to early September). Ten mature hazelnuts were collected from the ground under each of the trees that had previously been sampled for GSB individuals. To prevent them from rotting, the hazelnuts were dried in a ventilated container for two months and then stored in a refrigerated room at 13 °C. After two months, the nuts were shelled by hand one by one to extract the kernel, which was cut in half and visually inspected. White or brown necrosis spreading from the kernel’s surface to its center was attributed to GSB punctures (Hamidi *et al*., 2022).

### 1.3 Influence of adjacent habitats on GSB abundance in hazelnut orchards

#### Characterization of the adjacent habitats

The study was conducted at the five sites described in Section 1.2.

In front of each of the four opposite sides of a hazelnut orchard, an area measuring 100 x 50 meters was defined. These areas were located 5 meters from the edges of the orchard, directly in front of the rows that had been sampled for *P. prasina* abundance (see Section 1.2). A total of 32 areas corresponding to the three orchards monitored in 2021 and 2022, plus two additional orchards (one monitored exclusively in 2021, and one sampled only in 2022) were analyzed. The different types of habitats present in each area and in each year were mapped, and the relative area covered by each habitat was calculated using GIS tools (QGIS): the CES OSO layer (Thieron & Valero, 2016) and the BD Haie (IGN, 2023) were used in conjunction with photo interpretation (Taillefer, 1945). Due to the limited number of study areas (n = 32), the number of descriptive landscape variables was initially reduced by grouping some areas into broader habitat categories (Table 2): forest habitat (deciduous and coniferous forests, as well as hedges); grass-dominated habitat (grasslands, gardens, and grass strips); and habitats with impermeable surfaces (roads, scattered and dense buildings). An unscaled PCA was then applied to identify the primary variables, with minimal correlation, that contributed to variance in landscape composition among borders. Forest habitat, plum orchards, hazelnut orchards, and grassland areas were subsequently selected to characterize the landscape composition of each border.

**Table 2.** Characterization of habitat types

| Habitat types |  |  |  |
| --- | --- | --- | --- |
| Annual crops | Perennial crops | Forest habitats | Human structures |
| Cereals (Corn) | Hazelnut orchards | Deciduous forests | Roads |
| Protein crops (Soy, beans and other leguminous) | Plum orchards | Coniferous forests | Diffuse building |
| Sunflower | Wastelands | Hedges |  |
| Grasslands | Grass strips | - | Gardens |

#### Insect abundance in the hazelnut orchards

The data on the abundance of *P. prasina* in the orchards corresponds to the sampling results described in 1.2. For each side of the 4 sides of an orchard, the number of individuals collected in all rows during the full sampling period in the same year, were pooled together.

### 1.4 Statistical analyses

All the analyses described below were performed by fitting Generalized Linear Mixed Models (GLMMs) with the HLfit function from the spaMM package (version 4.4.0; Rousset & Ferdy, 2014) in R 4.3.1 (R Core Team, 2023). The GLMM analysis of GSB abundance data (pooled nymph and adult counts) used a Poisson distribution, while the GLMM analysis of damage used a Binomial distribution. For each model including interactions, an initial full model was first performed, which was then simplified by sequentially removing non-significant interaction terms to achieve the minimal adequate model. “Overdispersion” for each model was assessed by dividing the sum of the squared Pearson residuals by the residual degrees of freedom. This ratio should be less than 1 (Bolker *et al*., 2009). To test the significance of each fixed effect, we used the type II log likelihood ratio test (hereafter L.R.) to calculate P-values.

#### Host plant preferences

The “host preference” model was built using GSB abundance as the dependent variable. To keep things as simple as possible and to avoid collinearity of the independent variables, the two years (2021 and 2022) were analyzed separately, and phenological stage was first considered as presence or absence of fruits. We retained in the model the plant species (categorical; 10 plant species), phenological stage (categorical; 2 modalities), and their interaction as fixed effects. To account for spatial and temporal autocorrelation, we added two random effects on the intercept: the week date (autoregressive) and the plant ID (identifier) nested within the study site IDs.

#### GSB spatial distribution in hazelnut orchards

Due to the incomplete design in 2021, only the 2022 data was analyzed for the GSB spatial distribution. To account for the difference in the number of branches sampled per tree in old and young orchards and avoid biasing the orchard age effect, abundances on trees from young orchards were multiplied by two. To analyze the distribution of GSB at the beginning of orchard colonization, before farmers usually conduct pesticide treatments, the “population spatial distribution” model included the sum of GSB abundance over the first four weeks (20^th^ of June to 17^th^ of July - period 1) for each tree as dependent variable, with the orchard age (categorical; old orchard, and young orchards), the sampling distance (quantitative, with five distances 0, 5, 10, 15 and 30 meters) and their interactions as fixed effects. To keep the model as simple as possible, a ’tree_spacing’ variable with two levels (5 m and 2.5 m) was initially included in the first version of the model but was later removed as it was not significant and only served to control for heterogeneity. To account for spatial autocorrelation, the hazelnut tree ID nested within the four sides, themselves nested within the orchard ID, was added as a random effect. A second version of this same model was created using only the last 4 weeks of sampling (18^th^ of July to 14^th^ of August - period 2), to analyze the distribution of GSB at a later stage of orchards colonization. Again, the tree spacing effect was not significant and was therefore removed from the model. Using fixed-effects estimates from our models of GSB abundance during the first four weeks, we extrapolated predicted abundances from the orchard edge to its center (a distance of 150 meters), in 5-meter increments. We then calculated the cumulative relative abundance across this same range (0 to 150 m, in 5 m steps). To assess statistical uncertainty of this complex metric at each distance, we applied parametric bootstrap resampling (Kubokawa & Nagashima, 2011) with 300 replicates, enabling the construction of 95% confidence intervals. Finally, we identified the distance from the orchard edge within which at least 50% of the total GSB abundance was concentrated.

#### Damage spatial distribution in hazelnut orchards

The “damage spatial distribution” model included damage proportion (binary count; healthy or damaged) as the dependent variable, and the orchard age, sampling distance, and their interactions as fixed effects. To control for heterogeneity, a ’tree_spacing’ variable with two levels (5 m and 2.5 m) was included and was not subsequently removed because it was significant (Table 3). To account for spatial autocorrelation, the hazelnut tree ID nested within the four orchard sides, themselves nested within the orchard ID, was added as a random effect.

**Table 3.**
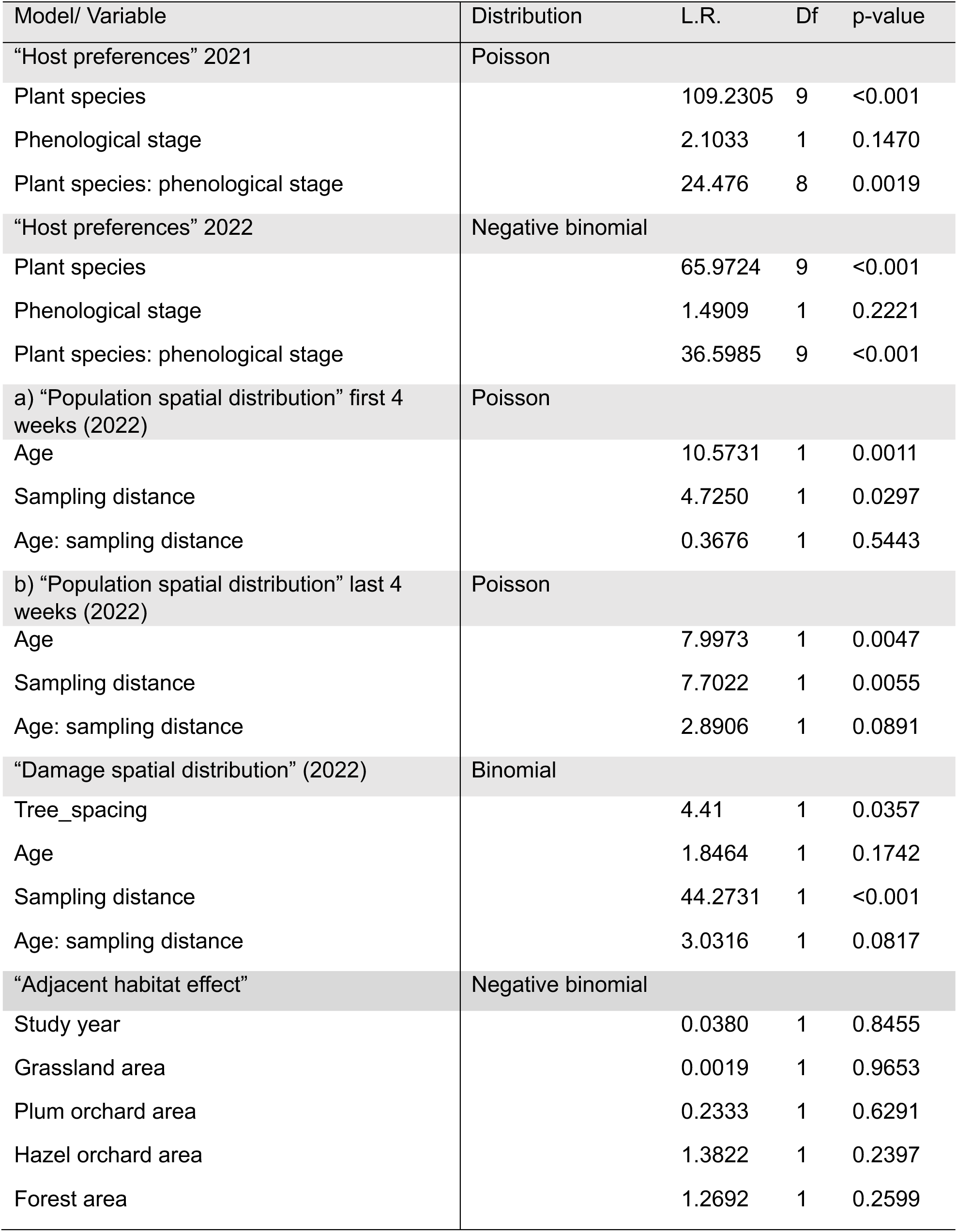
Results of the best fitting models (GLMM) used to test the effects of the plant species and adjacent habitats on the total abundance of GSB and to describe the spatial distribution of GSB population and damage proportion in hazelnut orchards.

#### Influence of adjacent habitats on GSB abundance in hazelnut orchards

The “adjacent habitat effect” model included GSB abundance pooled across combinations of orchards, sides, and years, as the dependent variable. The fixed predictor variables were the study year (continuous; 2021 and 2022) and the areas of grassland, plum orchards, hazelnut orchards, and forest habitats. The sample size was too small to add fixed interaction terms to the model. To account for spatial autocorrelation, we included orchard side nested within orchard ID as a random effect.

## Results

### 2.1 Host plant preferences

A total of 394 and 1144 individuals of GSB were captured in the hazelnut trees and adjacent hedge wild plants in 2021 and 2022, respectively; a total of 239 and 417 GSB individuals were collected from grass strips in 2021 and 2022, respectively.

Figures 1 and 2 show the fluctuation of the GSB population over time. Adults collected from April correspond to post-overwintering individuals (*i.e*. the first generation of adults). In both years, adults were found in hedge wild plants and in the hazelnut trees, but not in the grasses. They laid their first eggs from mid-April to the end of April, giving rise to the spring population. GSB nymphs corresponding to developmental stages 2 to 5 were collected in all sampled habitats (hazelnut trees, hedge wild plants, and grasses).

**Figure 1.**
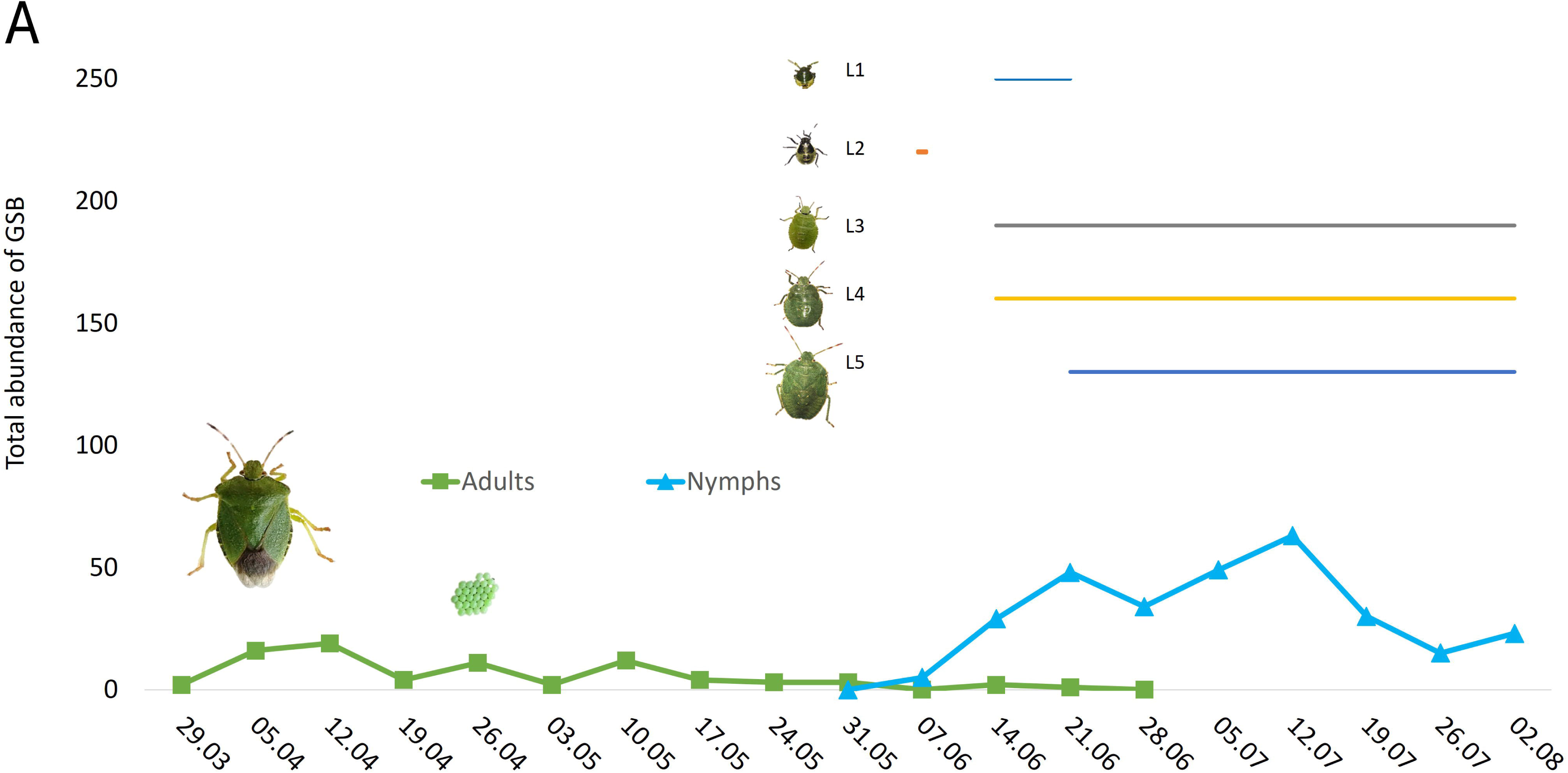
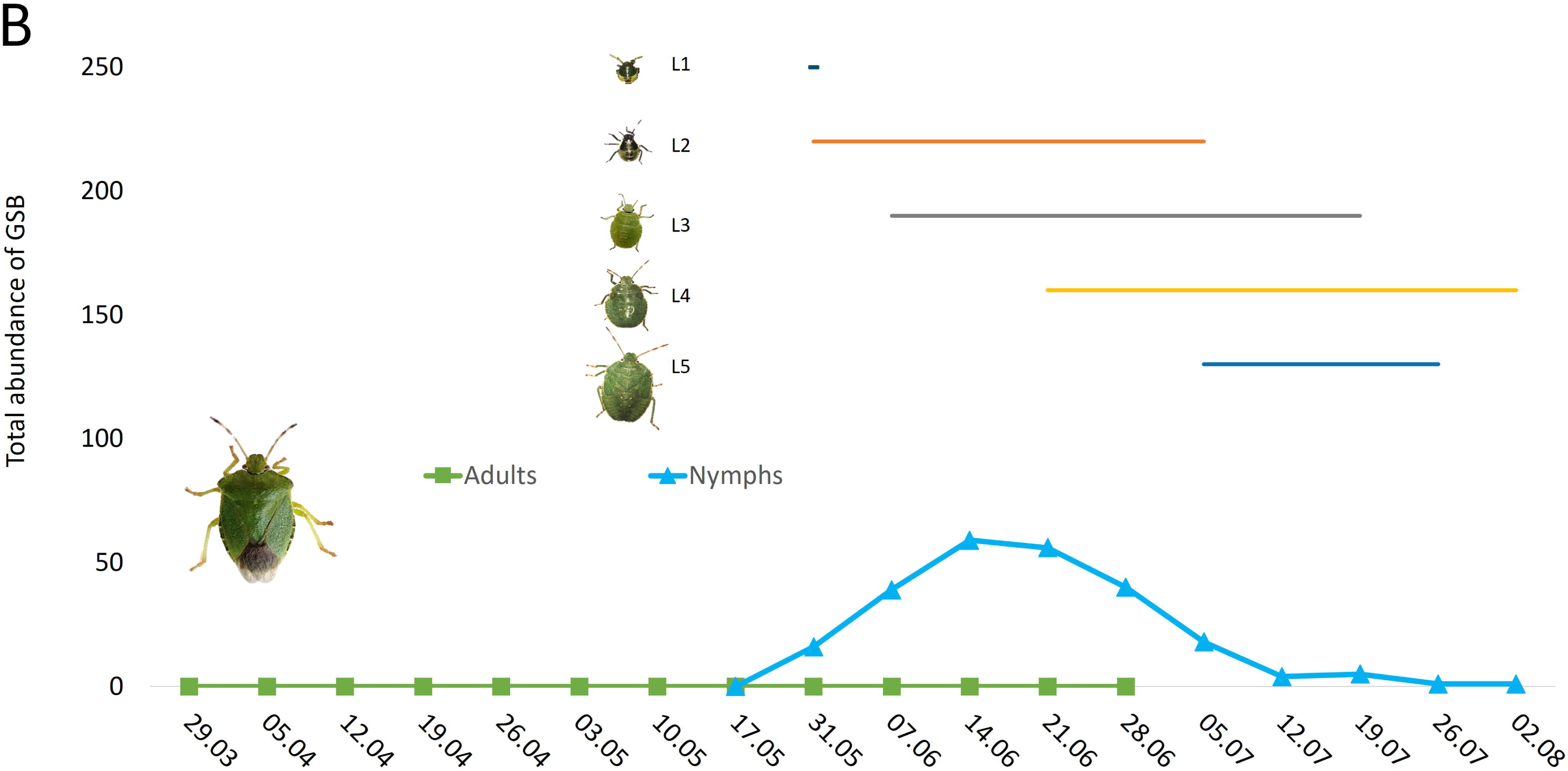
Monitoring of the seasonal variation in the abundance of *GSB* life stages in (A) woody plants (wild and hazelnut plants) and (B) herbaceous plants, in 2021. First spotted egg mass is indicated, and solid lines on the top right represent the periods of occurrence of the various nymphal stages of GSB.

**Figure 2.**
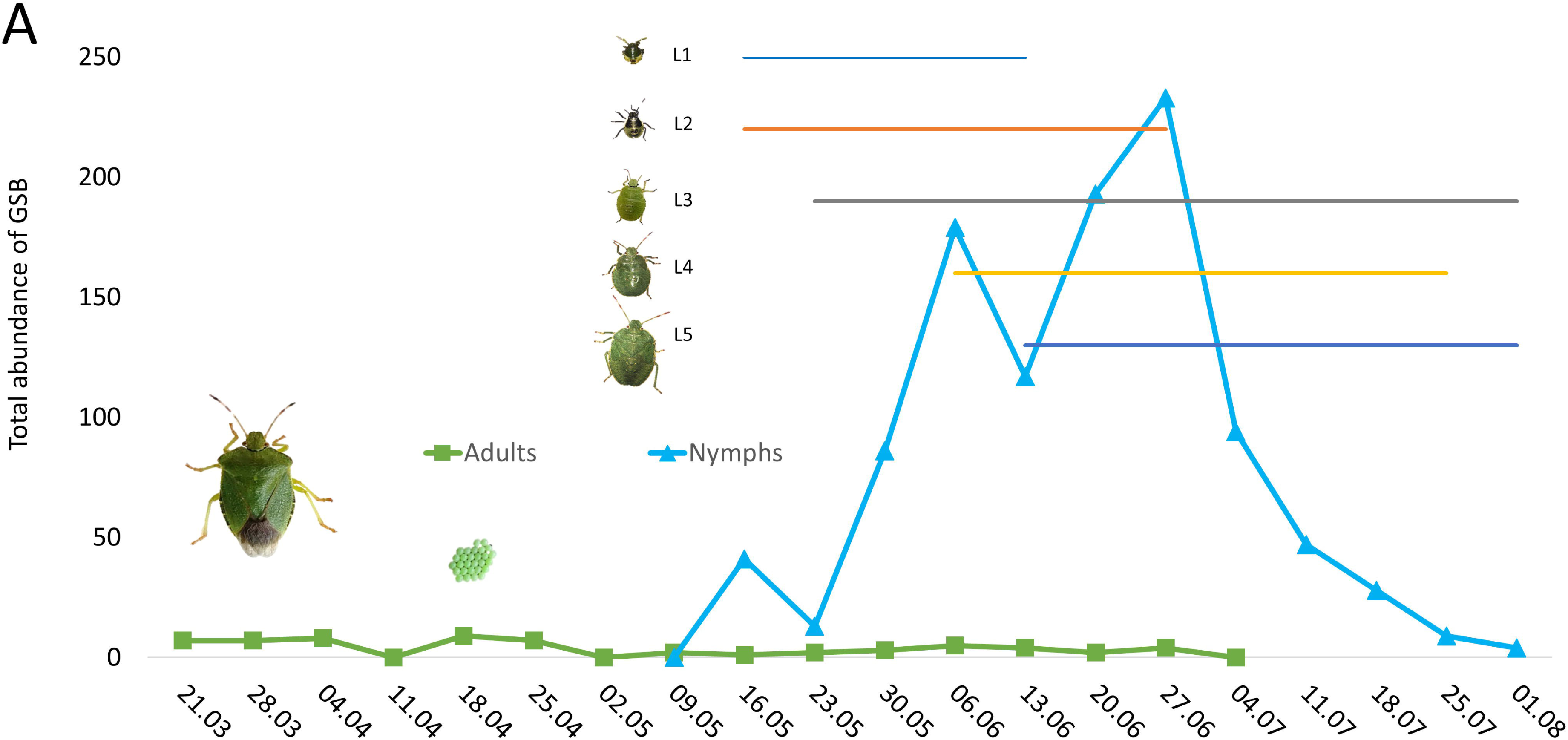
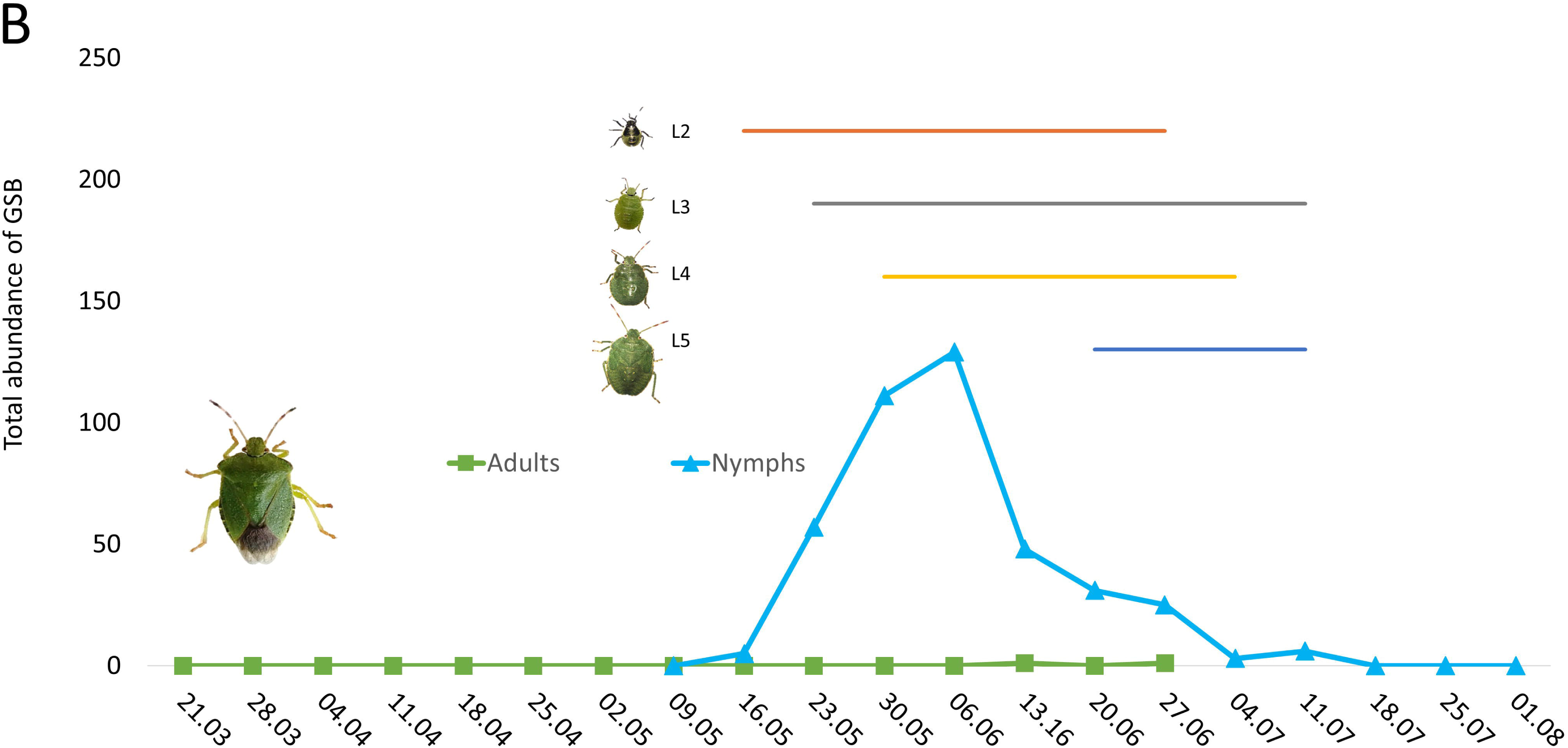
Monitoring of the seasonal variation in the abundance of *GSB* life stages in (A) woody plants (wild and hazelnut plants) and (B) herbaceous plants, in 2022. First spotted egg mass is indicated, and solid lines on the top right represent the periods of occurrence of the various nymphal stages of GSB.

Overall, GSB was present in all the plants studied, except on hazel trees in 2021. However, the abundance of GSB in the different woody plants on a same sampling date and over time varied (Supp. Mat. Table 1a and b). This could be related to plant species preference and plant phenological stage. The model testing for GSB host preferences shows a significant interaction between phenological stage and plant species for each year (Table 3; Figure 3), meaning that plant preference depends on the phenological stage considered. Nevertheless, a clear preference pattern emerges, with GSB being largely present in plants at the fruit stage and then the flower stage (Figure 3; Supp. Mat. Tables 1a and b). *C. sanguinea*, and *C. monogyna* were the plant species from which most individuals were consistently collected in 2021 and 2022 (Figure 3; Supp. Mat. Tables 1a and b). It should be noted that in 2022, when GSB was most abundant, *Rubus* sp. also joined this group of preferred plants. Unlike late instar nymphs, adults emerging from overwintering and 1^st^ and 2^nd^ instar nymphs do not colonize all plants.

**Figure 3.**
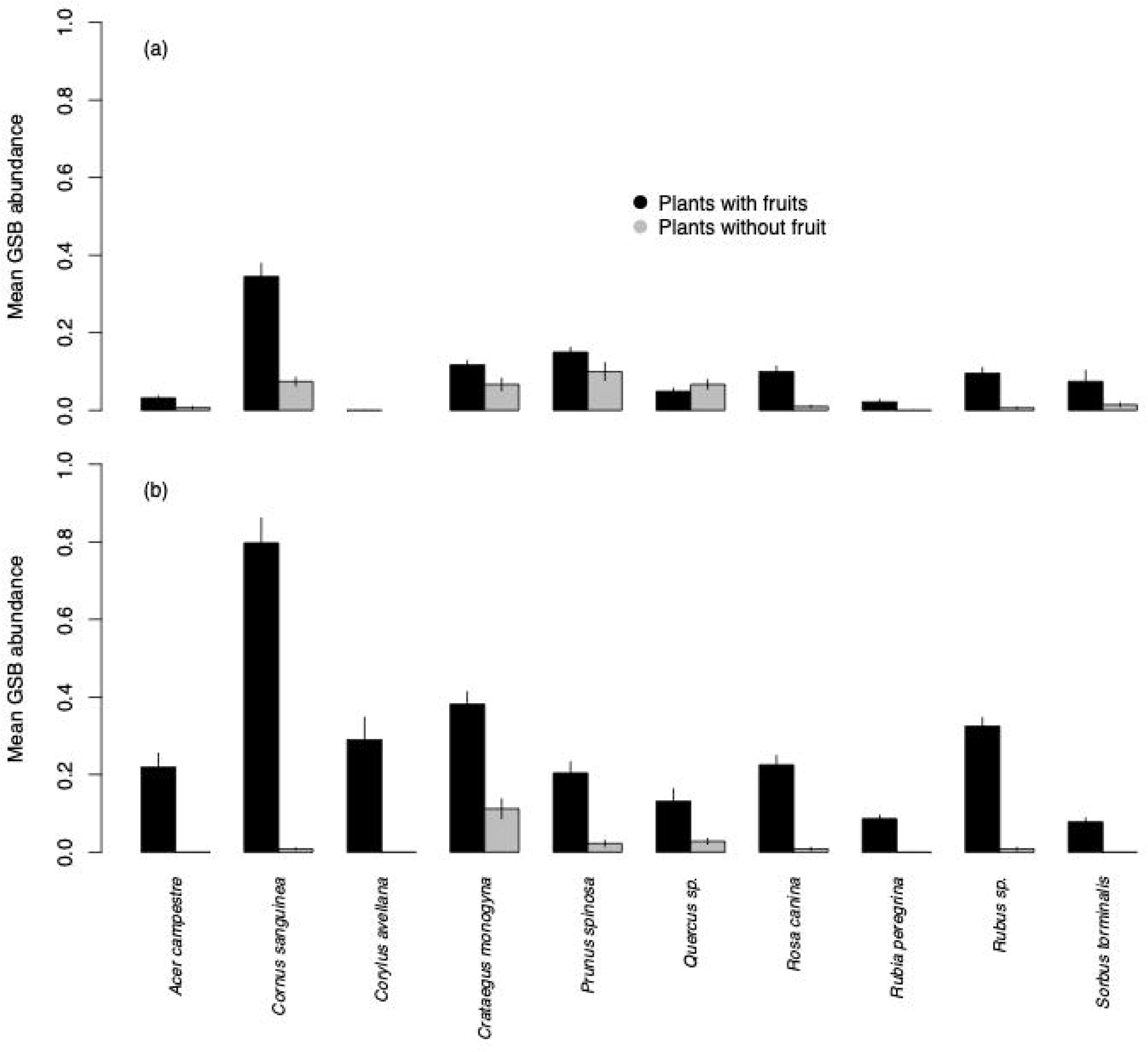
Mean (±SE) number of GSB individuals per tree with or without fruits for each plant species in 2021 (a) and 2022 (b).

### 2.2 GSB summer spatial distribution in hazelnut orchards

#### Distribution of GSB individuals

During the two-year study, a total of 663 GSB (312 in 2021 and 351 in 2022) were collected, accounting for 87.2 % of the Pentatomidae collected. They belonged exclusively to the 4^th^, 5^th^, and adult instars.

The model of the spatial distribution of GSB abundances in 2022, summed over the first four weeks, shows a significant difference in GSB abundance between old and young hazelnut orchards (Table 3), with a higher abundance in the former. Most importantly, there is a significant effect of the distance to the edge on the abundance of GSB, which decreases towards the center of the orchards (Figure 4a) with no interaction with age (Table 3). The same model applied to the data of the last 4 weeks of orchard colonization yielded identical results (Table 3). This time, the interaction between age and distance is marginally significant, with the effect of distance appearing to be stronger in younger orchards.

**Figure 4.**
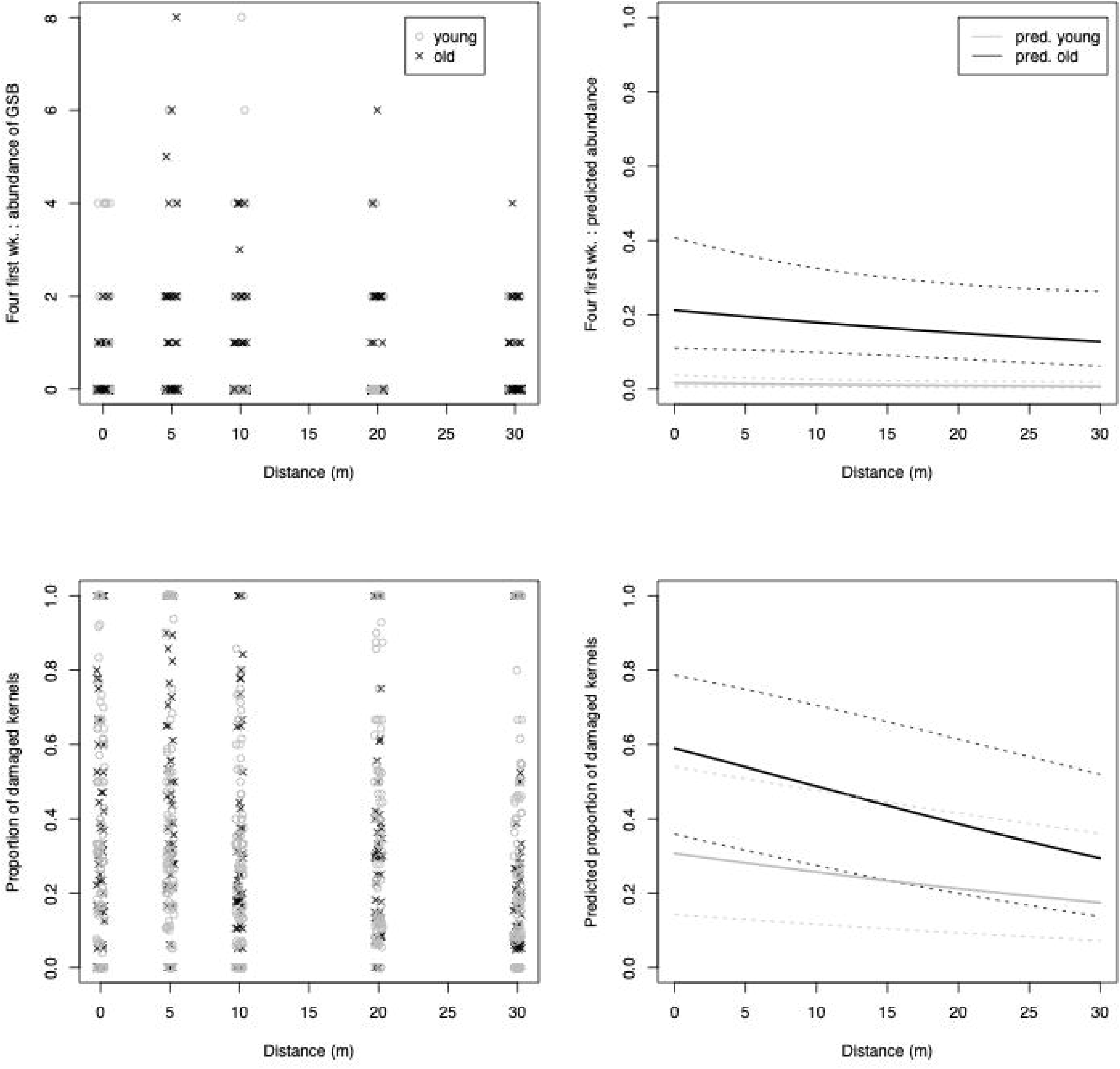
Spatial distribution in 2022 of a) GSB individuals during the first four weeks and b) damage proportion, at five different distances from the edge towards the centre of young and old hazelnut orchards. On the left, observed total abundance of GSB and damage proportion; on the right GLMM predictions (solid lines) with 95% confident intervals (dashed lines) taking into account variance due to uncertainty in fixed effects.

In both cases the relationship between GSB abundance and the distance to the edge is almost linear, suggesting that there are still some GSB beyond 30 meters to the center of the orchards. Based on the predicted total abundance of the first 4 weeks’ model and its parameters, we estimated that 50 % of the total GSB population would be within the first 25 meters from the edge (Figure 5). However, it should be noted that the confidence interval (CI) of the estimated distance threshold is relatively high.

**Figure 5.**
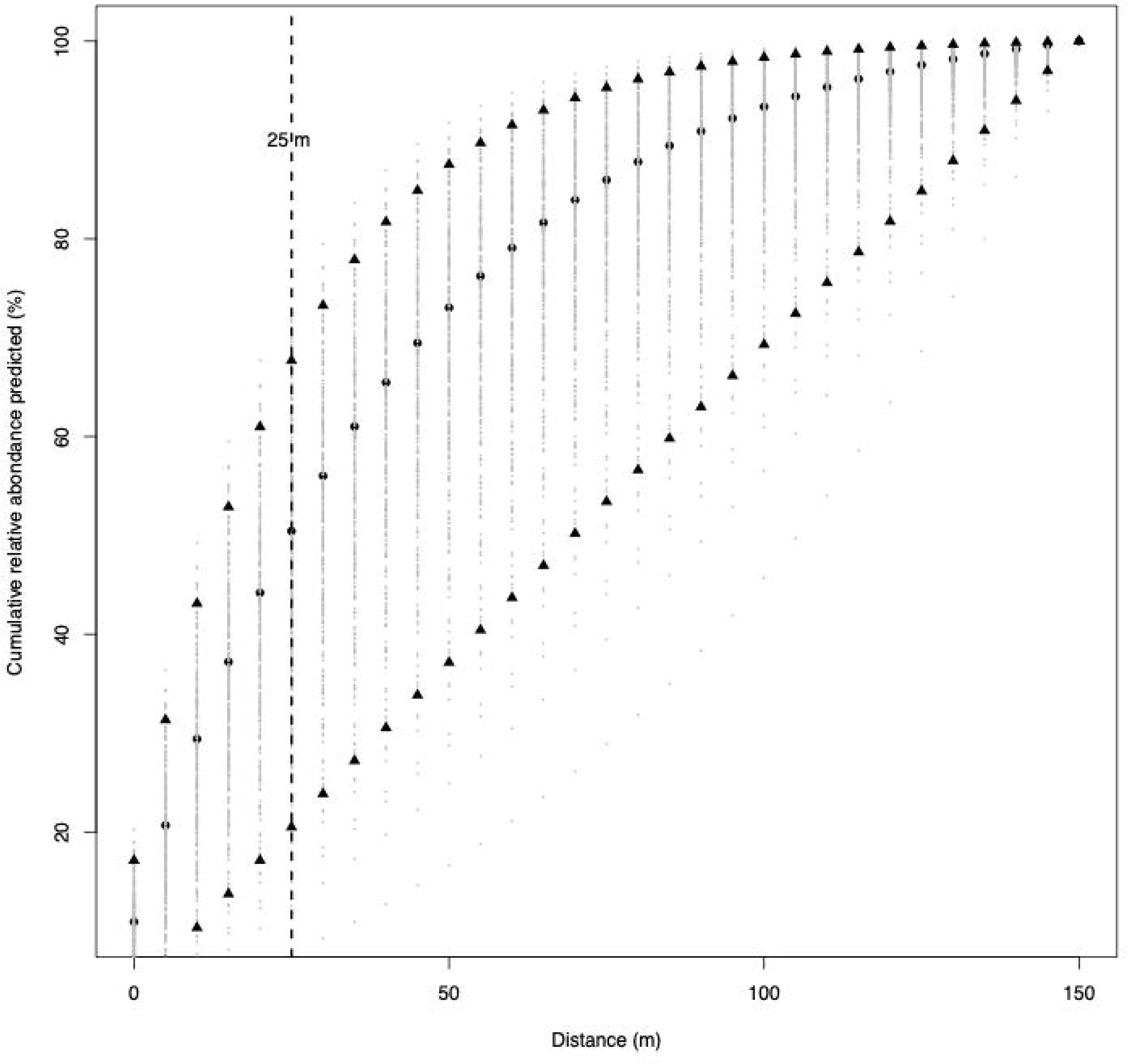
Predicted distance threshold where at least 50 % of GSB are collected. Dashed line = threshold; solid black dots= predicted abundance from the fitted model; solid grey dots= distribution of simulated bootstraps data, black triangles= upper and lower CI). This prediction was calculated using the partial model “population distribution” based on the first four weeks of sampling in 2022 and for old hazelnut orchard.

#### Distribution of damage

A total of 10597 hazelnuts were collected and analyzed, of which 948 were empty and 9 649 kernels were extracted. Seven per cent of the kernels were stunted or rotten and 1.4 % were partially eaten by the nut weevil, *Curculio nucum* L. (Coleoptera: Curculionidae); all these kernels were removed from the analysis.

Statistical analysis of the spatial distribution of damage shows a marginally significant interaction between orchard age and sampling distance. In both old and young orchards, the proportion of damaged hazelnuts decreases from the edge to the center, with a shallower gradient in young orchards (Table 3 and Figure 4b). Additionally, the proportion of GSB damage decreases more sharply from the edge to the center than GSB abundance (slope estimations for old orchards: −0.0412 and −0.0168, respectively).

2.3 Influence of adjacent habitats on GSB abundance in hazelnut orchards

None of the descriptive landscape variables were significant and can therefore be used to explain variation in GSB abundance in hazelnut orchards (Table 3). Nevertheless, it is noteworthy that the two *p-values* around 20 %, *i.e.* “hazelnut orchards” and “forest areas”, have a negative effect and positive effect on the GSB abundance, respectively (Table 3).

## Discussion

Currently, the control of the green shield bug (GSB) in hazelnut orchards across Europe mostly relies on the use of pesticides. Unfortunately, little research has been conducted into its ecology, despite this information being crucial for developing sustainable control measures. The main objective of this study was therefore to characterize the spatiotemporal dynamics of GSB populations from emergence from overwintering sites to the end of the crop production season, thereby paving the way for new IPM strategies for its control.

The monitoring of GSB on wild and hazelnut plants showed that it was present on all studied species throughout the sampling period, which is consistent with the polyphagous nature of stink bugs (Panizzi, 1997). Indeed, the development, fecundity and survival of species such as *Chinavia picticornis* (Stal), *H. halys,* and *Piezodorus guildinii* (Westwood), increase when they feed on a variety of wild and cultivated plants (Acebes-Doria *et al*., 2016b; da Silva *et al*., 2019; Zerbino *et al*., 2016). In addition, the nymphs of *Palomena angulosa* Motschulsky, a sister species of GSB, only complete their development when they feed on a variety of fruiting plants (Hori *et al*., 1985). Nevertheless, our study also indicates that, despite some differences between years, GSB prefers *C. monogyna* and *C. sanguinea,* where it has also been spotted in other countries (Bosco *et al*., 2017; Haye *et al*., 2020). Furthermore, our study reveals that although GSB is found in hazelnut plants (in 2022), it is only present there in late spring. *P. prasina* overwinters as adults in the leaf litter, particularly in hazelnut orchards, woods, and hedges (Driss *et al*., 2023). However, the fact that post-overwintering adults were found exclusively on wild plants, and not in cropped hazelnut trees, suggests that they move from the orchard trees to wild hosts in adjacent habitats in early spring, and only return later. Finally, our study shows that GSB is predominantly present in plants when they are at the fruit stage, which explains their later migration to hazelnut trees. As with many other Pentatomidae species, plants bearing fruits are the most attractive (Blaauw *et al*., 2019; Martinson *et al*., 2015). Adults and nymphs feed preferentially on fruiting structures, as these are the most nutrient-rich resources (Acebes-Doria *et al.,* 2016b; McPherson, 2018).

It should be noted that the present study of plant preferences is based on the relative abundance of individuals among different plants, making it an indirect measure of preference. Indeed, availability of resources may influence preference (Crawley, 1983) and, although we considered this by selecting the most common plant species in the region, we did not balance plant availability at each site. Moreover, the choice of an oviposition site by female insects is considered as a good indicator of the quality of the food source for the offspring (Resetarits, 1996), but we did not monitor eggs in our study. Although we have data on the presence of first-instar nymphs, whose movement is extremely limited and can therefore be used as a proxy for oviposition, the number of individuals at that developmental stage was very low. This makes it difficult to draw conclusions about plant preferences based on first-instar nymphs. Therefore, we consider that while our study establishes a foundation for investigating plant preferences in field conditions and therefore considers the strength of selection acting in the field, it should be complemented by behavioral and life-table experiments conducted in laboratory settings.

Our results show that, in spring, the highly mobile 3^rd^-5^th^ nymphal instars constitute the main dispersing population between the diverse host plants. Notably, they are also found in the herbaceous plants located between the hedge plants and the hazelnut orchards. These grass strips play a major role as a route in the subsequent dispersal of late-instar nymphs towards the orchards at the end of spring, which confirms Boselli’s (1932) earlier observations. The colonization of the orchards coincides with the fruitification of the hazelnut trees (Hamidi *et al*., 2022), which provide an abundant, high-quality, trophic resource. It is not uncommon for stink bug pests to move between wild plants and crops (Panizzi & Lucini, 2017). These seasonal fluctuations are primarily related to changes in the nutritional composition of hosts (Bakken *et al*., 2015). For example, late instar nymphs of *Euschistus servus* (Say) and *N. viridula* migrate from peanut to adjacent cotton crops when the latter attain an ideal nutritional quality (Tillman *et al*., 2009).

Movements between different habitats can lead to biased distributions and our results tend to confirm this, showing an edge-effect on the distribution of GSB in the hazelnut orchards. Indeed, we demonstrated that the abundance of the stink bug decreases from the edge towards the center of all the studied hazelnut orchards and that, most crucially, the damage proportion follows the same pattern. This corroborates the recent preliminary findings of Aymamí *et al*. (2023), who observed, for one orchard during a one-year sampling, that the abundance of GSB was higher at the edge than in the center of the orchard. Movement from wild plants to orchards, such as those observed in this study can lead to dilution effects towards the center of the orchard, which can be accentuated when the trees in the front rows are taller and/or more productive, as is often the case in temperate regions (Morreale *et al*., 2021). Due to their size, edge trees can act as barriers, restricting the movement of individuals - a phenomenon that has been observed for various stink bug species (Tillman, 2014). Furthermore, it has also been demonstrated that Pentatomidae nymphs, which constitute the main colonizing population of GSB, tend to move along rather than across crop rows (Bastola & Davis, 2018; Panizzi *et al*., 1980). This behavior could help explain the GSB edge-effect. Finally, Marshall and Beers (2024) have shown Pentatomidae movements back and forth between hedges and orchards throughout the season.

The presence of wooded habitats and hedges containing wild host plants, as well as certain crops, can lead to a higher abundance of stink bugs at the edges of adjacent crops (Tillman *et al*., 2009; Moraglio *et al*., 2018; Venugopal *et al*., 2014). Our study was unable to pinpoint a specific adjacent habitat type that may explain GSB abundance in hazelnut orchards during the production season. Nonetheless, GSB abundance tended to be positively associated with mixed forests and hedges, where GSB can be found feeding on various wild plant hosts in spring, and negatively associated with hazelnut orchards, despite the fact the orchards are overwintering sites. This is consistent with the aforementioned results showing that GSB leaves the orchards in early spring, only returning in late spring/early summer.

Based on all the above results, we hereafter suggest several methods for controlling *P. prasina* from an IPM perspective. Firstly, we established that GSB is polyphagous but prefers certain host plants. “Attract-and-kill” strategies (Shelton & Badenes-Perez, 2006) using preferred host plants as trap crops, have been employed to control Hemipteran pests. However, while the use of woody species is not ruled out (*e.g*., *Murraya paniculata* (L.) planted around orange groves attracts and kills 83 % of *Diaphorina citri* Kuwayama individuals (Tomaseto *et al*., 2019), herbaceous plants are easier to manage (Panizzi & Lucini, 2017). For instance, in the case of stink bugs, mustard has been successfully used in “Attract-and-kill” strategies against *N. viridula* (Wallingford *et al*., 2013). Our study also highlights the important role of annual plants in attracting GSB. Indeed, grass strips appear to be crucial for feeding and facilitating the dispersion of late stages towards the orchards. While Bulam Köse *et al*. (2014) tested the control of herbaceous weeds to reduce GSB populations, the authors emphasized the importance of knowing the exact moment to exert control. The importance of this timing of control was also noted by Babu *et al*. (2019) and Coombs (2000) when investigating the effect of destroying herbaceous plants next to crop fields on the populations of *N. viridula* and *E. servus.* Our study reveals that monitoring the colonization period in herbaceous plants is the most effective strategy for determining control timing in the case of GSB. Additionally, the use of obstacles such as plant-based or artificial barriers could restrict the spread of the colonizing GSB in orchards. This was demonstrated for certain species of stink bugs (e.g. Marshall & Beers, 2024; Tillman, 2014; Tillman *et al*., 2014).

Finally, we demonstrated the presence of an edge effect on GSB distribution in orchards. This edge-biased distribution could be important for decision making about the range of insecticide spraying. Indeed, our results show that in the early production season, when the first pesticide treatments are usually applied, almost 50 % of the GSB can be targeted at the edges. This could lead to a significant reduction in the amount of pesticides used. Edge targeting has been used to control *H. halys* (Herbert *et al*., 2015; Leskey *et al*., 2012), sometimes leading to a 2.5-fold reduction in pesticide treatments (Blaauw *et al*., 2015). However, it should be noted that our threshold estimation of 25 meters has some limitations, and further data is needed to improve the model by sampling beyond 30 meters.

In conclusion, we believe that this study, which highlighted various aspects of the spring-summer ecology of the GSB, provides valuable information for developing an efficient IPM strategy for controlling the GSB in hazelnut orchards.

## Acknowledgements

We would like to thank G. Richon and L. Chioso for assistance on GSB monitoring, and J. Robin, S. Papillon and A. Jagueneau, for help on damage assessment. This work was supported by Unicoque, the ANPN, and the European Community [FEDER FSE 2019 7838010 (REPLIK)], and we thank M. Thomas for contributing to founds acquisition. The laboratory “Centre de Recherche sur la Biodiversité et l’Environnement” is part of the “Laboratoire d’Excellence” LABEX TULIP (ANR-10-LABX-41) and LABEX CEBA (ANR-10-LABX-25-01). L. Driss benefited from a CIFRE PhD scholarship (CIFRE N°2020/1113)

## References

Acebes-Doria, A. L., Leskey, T. C., and Bergh, J. C. (2016a). Injury to apples and peaches at harvest from feeding by *Halyomorpha halys* (Stål) (Hemiptera: Pentatomidae) nymphs early and late in the season. Crop Protection, 89, 58–65. doi: 10.1016/j.cropro.2016.06.022

Acebes-Doria, A. L., Leskey, T. C. and Bergh, J. C. (2016b). Host plant effects on *Halyomorpha halys* (Hemiptera: Pentatomidae) nymphal development and survivorship. Environmental Entomology, 45(3), 663–670. doi: 10.1093/ee/nvw018

Aigner, B. L., Kuhar, T. P., Herbert, D. A., Brewster, C. C., Hogue, J. W. and Aigner, J. D. (2017). Brown marmorated stink bug (Hemiptera: Pentatomidae) infestations in tree borders and subsequent patterns of abundance in soybean fields. Journal of Economic Entomology, 110(2), 487–490. doi: 10.1093/jee/tox047

Ak, K., Tuncer, C., Baltacı, A., Eser, Ü. and Saruhan, İ. (2018). Incidence and severity of stink bugs damage on kernels in Turkish hazelnut orchards. Acta Horticulturae, 1226, 407–412. doi: 10.17660/ActaHortic.2018.1226.62

Aymamí, A., Barrios, G., Sordé, J., Palau, R., Solano, C., Domingo, L., Salvat, A. and Rovira, M. (2023). Biology and population dynamics of *Palomena prasina* L. in hazelnut orchards in Catalonia (Spain). Acta Horticulturae, 1379, 393–400. doi: 10.17660/ActaHortic.2023.1379.57

Babu, A., Reisig, D. D., Walgenbach, J. F., Heiniger, R. W. and Everman, W. (2019). Influence of weed manipulation in field borders on brown stink bug (Hemiptera: Pentatomidae) densities and damage in field corn. Environmental Entomology, 48(2), 444–453. doi : 10.1093/ee/nvz016

Badeau, V., Bonhomme, M., Bonne, F., Carré, J., Cecchini, S., Chuine, I., Ducatillion, C., Jean, F. and Lebourgeois, F. (2017). Les plantes au rythme des saisons : guide d’observation phénologique. Mèze, France : Biotope.

Bakken, A. J., Schoof, S. C., Bickerton, M., Kamminga, K. L., Jenrette, J. C., Malone, S., Abney, M. A., Herbert, D. A., Reisig, D., Kuhar, T. P. and Walgenbach, J. F. (2015). Occurrence of brown marmorated stink bug (Hemiptera: Pentatomidae) on wild hosts in non managed woodlands and soybean fields in North Carolina and Virginia. Environmental Entomology, 44(4), 1011–1021. doi: 10.1093/ee/nvv092

Bastola, A. and Davis, J. A. (2018). Determining in-field dispersal of the redbanded stink bug (Hemiptera: Pentatomidae) in soybean fields using a protein based mark-capture method. Crop Protection, 112, 24–32.

Bayle, M. (2022). Impact des attaques de punaises sur la productivité des vergers de noisetiers. Cancon, France: Unicoque.

Bergmann, E. J., Dilip Venugopal, P., Martinson, H. M., Raupp, M. J. and Shrewsbury, P. M. (2016). Host plant use by the invasive *Halyomorpha halys (Stål) o*n woody ornamental trees and shrubs. PLoS ONE, 11(2), 1–12.

Blaauw, B. R., Polk, D. and Nielsen, A. L. (2015). IPM-CPR for peaches: incorporating behaviorally-based methods to manage *Halyomorpha halys* and key pests in peach. Pest management science, 71(11), 1513–1522. doi: 10.1002/ps.3955

Bolker, B.M., Brooks, M.E., Clark, C.J., Geange, S.W., Poulsen, J.R., Stevens, M.H.H. and White, J.-S.S., (2009). Generalized linear mixed models: a practical guide for ecology and evolution. Trends in Ecology & Evolution 24, 127–135. 10.1016/j.tree.2008.10.008

Bosco, L., Moraglio, S. T. and Tavella, L. (2017). *Halyomorpha halys*, a serious threat for hazelnut in newly invaded areas. Journal of Pest Science, 91, 661–670. doi: 10.1007/s10340-017-0937-x

Boselli, F.B. (1932). Studio biologico degli Emitteri che attaccano le nocciuole in Sicilia. *Rivista di Patologia Vegetale*, Serie II, 22(9/10) : 333-335.

Bulam Köse, Ç., Sezer, A. and IġIK, D. (2014) Farklı yabancı ot mücadele yöntemlerinin Fındık Kokarcası [(*Palomena prasina* L.) (Hemiptera: Pentatomidae)] popülasyonu ve zarar durumuna etkisinin belirlenmesi. Bitki Koruma Bülteni, 54(*1*),79–92

Chocorosqui, V. R. and Panizzi, A. R. (2004). Impact of cultivation systems on *Dichelops melacanthus* (Dallas) (Heteroptera: Pentatomidae) population and damage and its chemical control on wheat. Neotropical Entomology, 33(4), 487–492. doi: 10.1590/S1519-566X2004000400014

Cissel, W. J., Mason, C. E., Whalen, J., Hough-Goldstein, J. and Hooks, C. R. R. (2015). Effects of Brown Marmorated Stink Bug (Hemiptera: Pentatomidae) Feeding Injury on sweet corn yield and quality. Journal of Economic Entomology, 108(3), 1065–1071. doi: 10.1093/jee/tov059

Coombs, M. T. (2000). Seasonal phenology, parasitism, and evaluation of mowing as a control measure for *Nezara viridula* (Hemiptera: Pentatomidae) in Australian pecans. Environmental Entomology, 29(5), 1027–1033. doi: 10.1603/0046-225X-29.5.1027

Crawley, M. J. (1983). Herbivory. The dynamics of animal-plant interactions. Oxford, United Kingdoms: Blackwell Scientific Publications.

da Silva, C. C. A., Blassioli-Moraes, M. C., Borges, M. and Laumann, R. A. (2019). Food diversification with associated plants increases the performance of the Neotropical stink bug, *Chinavia impicticornis* (Hemiptera: Pentatomidae). Arthropod-Plant Interactions, 13(3), 423–429. doi: 10.1007/s11829-018-9637-6

Driss, L., Hamidi, R., Andalo, C. and Magro, A. (2023). Study of the overwintering ecology of the hazelnut pest, *Palomena prasina* (L.) (Hemiptera: Pentatomidae) in a perspective of Integrated Pest Management. Journal of Applied Entomology, 148(*1*),34–48. doi: 10.1111/jen.13206

Esquivel, J. F., Musolin, D. L., Jones, W. A., Rabitsch, W., Greene, J. K., Toews, M. D., Schwertner, C., Grazia, J. and McPherson, J. E. (2018). Nezara viridula (L.). In Invasive stink bugs and related species (Pentatomoidea): biology, higher systematics, semiochemistry, and management (Mc Pherson, pp. 351–423). CRC Press.

Gerson, U. and Cohen, E. (1989). Resurgences of spider mites (Acari: Tetranychidae) induced by synthetic pyrethroids. Experimental and Applied Acarology, 6(1), 29–46. doi: 10.1007/BF01193231

Grazia, J., Panizzi, A., Greve, C., Schwertner, C., Campos, L., Garbelotto, T. and Fernandes, J. (2015). Stink Bugs (Pentatomidae), In A. R. Panizzi & J. Grazia (Eds.) True Bugs (Heteroptera) of the Neotropics (pp. 681–756). Dordrecht, The Netherlands: Springer Netherlands. doi: 10.1007/978-94-017-9861-7_22

Hamidi, R., Rouzes, R., Toillon, J., Thomas, M. and Tavella, L. (2023). The green shield bug, *Palomena prasina*, and the red-legged shield bug, *Pentatoma rufipes*, two secondary pests of French hazelnuts. Acta Horticulturae, 1379, 385–392. doi: 10.17660/ActaHortic.2023.1379.56

Hamidi, Rachid, Calvy, M., Valentie, E., Driss, L., Guignet, J., Thomas, M. and Tavella, L. (2022). Symptoms resulting from the feeding of true bugs on growing hazelnuts. Entomologia Experimentalis et Applicata, 170(6), 477–487. doi: 10.1111/eea.13165

Haye, T., Moraglio, S. T., Stahl, J., Visentin, S., Gregorio, T. and Tavella, L. (2020). Fundamental host range of *Trissolcus japonicus* in Europe. Journal of Pest Science, 93, 171–182. doi: 10.1007/s10340-019-01127-3

Hedstrom, C. S., Shearer, P. W., Miller, J. C. and Walton, V. M. (2014). The effects of kernel feeding by *Halyomorpha halys* (Hemiptera: Pentatomidae) on commercial hazelnuts. Journal of Economic Entomology, 107(5), 1858–1865. doi: 10.1603/EC14263

Herbert, D. A., Cissel, W. J., Whalen, J., Dively, G. and Hooks, C. R. R. (2015). *Brown marmorated stink bug biology and management in mid-Atlantic soybeans*. ENTO-168NP. Va. Coop. Ext

Hori, K., Kuramochi, K. and Nakabayashi, S. (1985). Effect of several different food plants on nymphal development of *Palomena angulosa* Motschulsky (Hemiptera: Pentatomidae). Research Bulletin of Obihiro University,14, 239–246.

Huang, T.-I., Reed, D. A., Perring, T. M. and Palumbo, J. C. (2014). Feeding damage by *Bagrada hilaris* (Hemiptera: Pentatomidae) and impact on growth and chlorophyll content of Brassicaceous plant species. Arthropod-Plant Interactions, 8(2), 89–100. doi: 10.1007/s11829-014-9289-0

IGN (2023). BD Haie. https://geoservices.ign.fr/bdhaie. Accessed 17 October 2023.

Iost Filho, F. H., Heldens, W. B., Kong, Z. and de Lange, E. S. (2020). Drones: innovative technology for use in precision pest management. Journal of Economic Entomology, 113(1), 1–25. doi: 10.1093/jee/toz268

Javahery, M. (1967). The biology of some Pentatomidae and their egg parasites. Faculty of Science of the University of London.

Kathage, J., Castañera, P., Alonso-Prados, J. L., Gómez-Barbero, M. and Rodríguez-Cerezo, E. (2018). The impact of restrictions on neonicotinoid and fipronil insecticides on pest management in maize, oilseed rape and sunflower in eight European Union regions. Pest management science, 74(1), 88–99. doi: 10.1002/ps.4715

Kubokawa, T. and Nagashima, B. (2011). Parametric bootstrap methods for bias correction in linear mixed models. J. Multivariate Anal. 106: 1–16. doi10.1016/j.jmva.2011.12.002

Leskey, T. C., Hamilton, G. C., Nielsen, A. L., Polk, D. F., Rodriguez-Saona, C., Bergh, J. C. et al. (2012). Pest status of the brown marmorated stink bug, Halyomorpha halys in the USA. Outlooks on Pest Management, 23(5), 218–226. doi: 10.1564/23oct07

Lupoli, R. and Dusoulier, F. (2015). Les punaises Pentatomoidea de France. Fontenay-sous-Bois, France: Éditions Ancyrosoma.

Marshall, A. T. and Beers, E. H. (2024). Using stink bug migration behavior for physical exclusion. Environmental Entomology, 53(3), 338–346. doi: 10.1093/ee/nvae025

Martinson, H. M., Venugopal, P. D., Bergmann, E. J., Shrewsbury, P. M. and Raupp, M. J. (2015). Fruit availability influences the seasonal abundance of invasive stink bugs in ornamental tree nurseries. Journal of Pest Science, 88: 461–468. doi10.1007/s10340-015-0677-8

McPherson, J.E. (2018). Invasive stink bugs and related species (Pentatomoidea): biology, higher systematics, semiochemistry, and management. CRC Press : 842 pp.

McPherson, J. E. and McPherson, R. (2000). Stink bugs of economic importance in America north of Mexico. CRC Press : 272 pp.

Miranda, M. P., Eduardo, W. I., Tomaseto, A. F., Volpe, H. X. L. and Bachmann, L. (2021). Frequency of processed kaolin application to prevent *DIAPHORINA CITRI* infestation and dispersal in flushing citrus orchards. Pest Management Science, 77(12), 5396–5406. doi: 10.1002/ps.6579

Moraglio, S. T., Bosco, L. and Tavella, L. (2018). *Halyomorpha halys* invasion: a new issue for hazelnut crop in northwestern Italy and western Georgia? Acta Horticulturae, 1226, 379–384. doi: 10.17660/ActaHortic.2018.1226.58

Morreale, L. L., Thompson, J. R., Tang, X., Reinmann, A. B. and Hutyra, L. R. (2021). Elevated growth and biomass along temperate forest edges. Nature Communications, 12(1), 7181. doi: 10.1038/s41467-021-27373-7

Nguyen, H. D. D., and Nansen, C. (2018). Edge-biased distributions of insects. A review. Agronomy for Sustainable Development, 38(1), 11. doi: 10.1007/s13593-018-0488-4

Panizzi, A. R. (1997). Wild hosts of Pentatomids: Ecological significance and role in their pest status on crops. Annual Review of Entomology, 42(1), 99–122. doi: 10.1146/annurev.ento.42.1.99

Panizzi, A. R., Galileo, M. H., Gastal, H. A., Toledo, J. F. and Wild, C. H. (1980). Dispersal of *Nezara viridula* and *Piezodorus guildinii* nymphs in soybeans. Environmental Entomology, 9(3), 293–297. 10.1093/ee/9.3.293

Panizzi, A. and Lucini, T. (2017). Host Plant-Stink Bug (Pentatomidae) Relationships. In A. Čokl & M. Borges (Eds.), Stink Bugs (pp. 31–58). Boca Raton, United States: CRC Press. doi: 10.1201/9781315120713-3

R Core Team. (2023). R: A language and environment for statistical computing. Vienna: R Foundation for Statistical Computing. Retrieved from https://www.R-project.org/

Ravula, A. R. and Yenugu, S. (2021). Pyrethroid based pesticides–chemical and biological aspects. Critical Reviews in Toxicology, 51(2), 117–140.

Resetarits Jr, W. J. (1996). Oviposition site choice and life history evolution. American Zoologist, 36(2) : 205–215.

Rice, K. B., Bergh, C. J., Bergmann, E. J., Biddinger, D. J., Dieckhoff, C., Dively, G. et al. (2014). Biology, ecology, and management of brown marmorated stink bug (Hemiptera: Pentatomidae). Journal of Integrated Pest Management, 5(3), A1–A13. doi: 10.1603/IPM14002

Romero, A., Tous, J. and Martí, E. (2009). White spots in hazelnut kernel: symptoms, causes and quality loss. Acta Horticulturae, 845, 607–612. doi: 10.17660/ActaHortic.2009.845.95

Rousset, F. and Ferdy, J.-B. (2014). Testing environmental and genetic effects in the presence of spatial autocorrelation. Ecography, 37, 781–790. 10.1111/ecog.00566

Saruhan, İ., Tunçer, M. K. and Tuncer, C. (2023). Economic damage levels of the green shield bug (*Palomena prasina*, Hemiptera: Pentatomidae) in Türkiye hazelnut orchards. Black Sea Journal of Agriculture, 6 (2): 183–189.

Sétamou, M. and Bartels, D. W. (2015). Living on the edges: spatial niche occupation of asian citrus psyllid, *Diaphorina citri* Kuwayama (Hemiptera: Liviidae), in citrus groves. PLOS ONE, 10(7), e0131917. doi: 10.1371/journal.pone.0131917

Shelton, A. M. and Badenes-Perez, F. R. (2006). Concepts and applications of trap cropping in pest management. Annual review of entomology, 51(1), 285–308.

Sosa-Gómez, D. R., Corrêa-Ferreira, B. S., Kraemer, B., Pasini, A., Husch, P. E., Delfino Vieira, C. E., et al. (2020). Prevalence, damage, management and insecticide resistance of stink bug populations (Hemiptera: Pentatomidae) in commodity crops. Agricultural and Forest Entomology, 22(2), 99–118. doi: 10.1111/afe.12366

Taillefer, F. (1945). La photographie aérienne au service de la géographie du Sud-Ouest. Revue géographique des Pyrénées et du Sud-Ouest. Sud-Ouest Européen, 16 (1): 143–150.

Tavella, L., Arzone, A., Miaja, M. L. and Sonnati, C. (2001). Influence of bug (Heteroptera, Coreidae and Pentatomidae) feeding activity on hazelnut in northwestern Italy. Acta Horticulturae, 556, 461–468. doi: 10.17660/ActaHortic.2001.556.68

Thieron V and Valero S. (2016). Carte d’occupations des sols de la France métropolitaine. https://www.theia-land.fr/product/carte-doccupation-des-sols-de-la-france-metropolitaine/. Accessed 17 October 2023.

Tillman, P. G., Northfield, T. D., Mizell, R. F. and Riddle, T. C. (2009). Spatiotemporal patterns and dispersal of stink bugs (Heteroptera: Pentatomidae) in peanut-cotton farmscapes. Environmental Entomology, 4(38), 1038–1052.

Tillman, P. G. (2014). Physical barriers for suppression of movement of adult stink bugs into cotton. Journal of Pest Science, 87(3), 419–427. doi: 10.1007/s10340-014-0564-8

Tillman, P.G., Cottrell, T. E., Mizell, R. F. and Kramer, E. (2014). Effect of field edges on dispersal and distribution of colonizing stink bugs across farmscapes of the Southeast USA. Bulletin of Entomological Research, 104(1), 56–64. doi: 10.1017/S0007485313000497

Tomaseto, A. F., Marques, R. N., Fereres, A., Zanardi, O. Z., Volpe, H. X., Alquezar, B., et al. (2019). Orange jasmine as a trap crop to control *Diaphorina citri*. Scientific Reports, 9(1), 2070.

Tuncer, C., Saruhan, I. and Akça, I. (2005). The insect pest problem affecting hazelnut kernel quality in Turkey. Acta Horticulturae, 686, 367–376. doi: 10.17660/ActaHortic.2005.686.51

Venugopal, P. D., Coffey, P. L., Dively, G. P. and Lamp, W. O. (2014). Adjacent habitat influence on stink bug (Hemiptera: Pentatomidae) densities and the associated damage at field corn and soybean edges. PLoS ONE ,10(9), Article 9.

Vernon, R. and Van Herk, W. (2017). Wireworm and flea beetle IPM in potatoes in Canada: implications for managing emergent problems in Europe. Potato Research, 60(3–4), 269–285. doi: 10.1007/s11540-018-9355-6

Wallingford, A. K., Kuhar, T. P., Pfeiffer, D. G., Tholl, D. B., Freeman, J. H., Doughty, H. B. and Schultz, P. B. (2013). Host plant preference of harlequin bug (Hemiptera: Pentatomidae), and evaluation of a trap cropping strategy for its control in collard. Journal of Economic Entomology, 106(1), 283–288. doi: 10.1603/EC12214

Wiman, N., Pscheidt, J. W. and Moretti, M. (2023). *Hazelnut Pest Management Guide for the Willamette Valley*. Retrieved from https://extension.oregonstate.edu/catalog/pub/em8328

Zerbino, M. S., Altier, N. A. and Panizzi, A. R. (2016). Performance of nymph and adult of *Piezodorus guildinii* (Westwood) (Hemiptera: Pentatomidae) feeding on cultivated legumes. Neotropical Entomology, 45(2), 114–122.

