## Supplementary material for "Edge effect on the distribution of the Green Shield Bug *Palomena prasina* in hazelnut orchards, and the role of adjacent habitats in crop colonization": Suplemental table 1

Supplementary material Table 1a. Relative abundance (%) of *P. prasina* monitored in wild hedge plant species and in hazelnuts from March to August 2021. N= Total absolute abundance. Vegetative period, Flower period, Fruit period

| **Plant species** | **March** | **April** | | | | **May** | | | | **June** | | | | **July** | | | | | **August** |
|---|---|---|---|---|---|---|---|---|---|---|---|---|---|---|---|---|---|---|---|
| | 29/03 | 05/04 | 12/04 | 19/04 | 26/04 | 03/05 | 10/05 | 17/05 | 25/05 | 01/06 | 07/06 | 14/06 | 21/06 | 01/07 | 05/07 | 12/07 | 19/07 | 26/07 | 02/08 |
| *A. campestre* | | | 5.6 | | | 50.0 | | | | | | 3.2 | | 2.9 | 6.4 | 4.5 | 12.9 | | |
| *C. sanguinea* | 50.0 | 26.7 | 5.6 | | 45.5 | | 58.3 | 25.0 | 12.5 | 66.7 | | 12.9 | 52.1 | 44.1 | 34.0 | 28.4 | 19.4 | 16.7 | 20.0 |
| *C. avellana* | | | | | | | | | | | | | | | | | | | |
| *C. monogyna* | | 6.7 | 38.9 | 25.0 | 9.1 | | 8.3 | 50.0 | | | | 3.2 | 14.6 | 5.9 | 10.6 | 25.4 | 25.8 | 8.3 | 16.0 |
| *P. spinosa* | | 40.0 | 22.2 | 25.0 | 27.3 | | 33.3 | 25.0 | 12.5 | 8.3 | 40.0 | 3.2 | 12.5 | 17.6 | 21.3 | 25.4 | 25.8 | 33.3 | 24.0 |
| *Quercus* sp. | 50.0 | 13.3 | 27.8 | | 18.2 | 50.0 | | | 62.5 | 8.3 | 20.0 | 3.2 | | 5.9 | 4.3 | 6.0 | 3.2 | | 8.0 |
| *R. canina* | | 6.7 | | | | | | | 12.5 | 8.3 | | 6.5 | 6.3 | 20.6 | 12.8 | 3.0 | 3.2 | | 20.0 |
| *Rubus* sp. | | | | | | | | | | | 40.0 | 3.2 | 14.6 | 2.9 | 6.4 | 6.0 | 9.7 | 25.0 | |
| *R. peregrina* | | | | | | | | | | | | | | | | 1.5 | | 8.3 | 4.0 |
| *S. torminalis* | | 6.7 | | 50.0 | | | | | | 8.3 | | 64.5 | | | 4.3 | | | 8.3 | 8.0 |
| N | 2 | 15 | 18 | 4 | 11 | 2 | 12 | 4 | 8 | 12 | 5 | 31 | 48 | 34 | 47 | 67 | 31 | 12 | 25 |

Supplementary material Table 1b. Relative abundance (%) of *P. prasina* monitored in wild hedge plant species and in hazelnuts from March to August 2022. N= Total absolute abundance. Vegetative period, Flower period, Fruit period

| Plant species | March | | April | | | | May | | | | June | | | | July | | | | | August |
|---|---|---|---|---|---|---|---|---|---|---|---|---|---|---|---|---|---|---|---|---|
| | 21/03 | 28/03 | 04/04 | 11/04 | 18/04 | 25/04 | 02/05 | 09/05 | 16/05 | 24/05 | 01/06 | 06/06 | 13/06 | 20/06 | 27/06 | 04/07 | 11/07 | 18/07 | 27/07 | 01/08 |
| *A. campestre* | | | | | | | | | 58.5 | 21.4 | 1.1 | 15.5 | 5.0 | 4.7 | 2.6 | 9.6 | | 25.0 | 23.5 | 55.6 |
| *C. sanguinea* | | | | | | | | 100.0 | | | 1.1 | 8.3 | 36.7 | 46.8 | 35.9 | 26.0 | 39.3 | 11.1 | 5.9 | 11.1 |
| *C. avellana* | | | | | | | | | 2.4 | | 46.6 | 22.1 | 19.2 | 1.6 | | 1.9 | 3.6 | 11.1 | | |
| *C. monogyna* | 71.4 | | 100.0 | | 44.4 | 28.6 | | | 36.6 | | 10.2 | 0.6 | 6.7 | 15.3 | 23.4 | 20.2 | 21.4 | 16.7 | 41.2 | 22.2 |
| *P. spinosa* | 28.6 | | | | 11.1 | | | | | | 1.1 | 18.2 | 4.2 | 9.5 | 9.1 | 14.4 | 3.6 | 16.7 | 11.8 | |
| *Quercus* sp. | | | | | 22.2 | 42.9 | | | | | 6.8 | 14.4 | 2.5 | 2.1 | 1.7 | 5.8 | 7.1 | 2.8 | 5.9 | |
| *R. canina* | | | | | | 28.6 | | | | 14.3 | 2.3 | 5.5 | 3.3 | 10.0 | 11.3 | 9.6 | 5.4 | 8.3 | 11.8 | |
| *Rubus* sp. | | | | | 22.2 | | | | 2.4 | 57.1 | 26.1 | 9.9 | 13.3 | 6.3 | 10.8 | 8.7 | 8.9 | | | |
| *R. peregrina* | | | | | | | | | | | 2.3 | 3.3 | 7.5 | 1.6 | 1.3 | 3.8 | 7.1 | | | |
| *S. torminalis* | | | | | | | | | | 7.1 | 2.3 | 2.2 | 1.7 | 2.1 | 3.9 | | 3.6 | 8.3 | | 11.1 |
| N | 7 | 0 | 8 | 0 | 9 | 7 | 0 | 2 | 41 | 14 | 88 | 181 | 120 | 190 | 231 | 104 | 56 | 36 | 17 | 9 |
